# Structural and dynamical heterogeneity of water trapped inside Na^+^-pumping KR2 rhodopsin in the dark state

**DOI:** 10.1101/2020.09.28.316596

**Authors:** Mantu Santra, Aniruddha Seal, Kankana Bhattacharjee, Suman Chakrabarty

## Abstract

Photoisomerisation in retinal leads to a channel opening in the rhodopsins that triggers translocation or pumping of an ion/proton. Crystal structures of rhodopsins contain several structurally conserved water molecules. It has been suggested that water plays an active role in facilitating the ion pumping/translocation process by acting as a lubricant in these systems. In this work, we systematically investigate the localisation, structure, dynamics and energetics of the water molecules along the channel for the resting/dark state of KR2 rhodopsin. Employing several microseconds long atomistic molecular dynamics (MD) simulation of this trans-membrane protein system, we demonstrate the presence of five distinct water containing pockets/cavities separated by gateways controlled by the protein side-chains. There exists a strong hydrogen bonded network involving these buried water molecules and functionally important key residues. We present evidence of significant structural and dynamical heterogeneity in the water molecules present in these cavities with very rare exchange between them. The exchange time-scale of these buried water with bulk has an extremely wide range from tens of nanoseconds to > 1.5*μs*! The translational and rotational dynamics of buried water are found to be strongly dependent on the protein cavity size and local interactions with classic signature of trapped diffusion and rotational anisotropy.

## Introduction

The structure, dynamics and function of biomolecules such as proteins and DNA heavily depend on their solvation environment. In this context, extensive studies have been carried out to investigate the active role of “biological water”.^1–12^ Hydration level and water dynamics near active sites often strongly influence the functionality of proteins and protein-DNA complexes.^13–15^ The dynamics of water at the surface or in the interior of proteins can be coupled to the protein dynamics and can occur at a wide range of time-scales depending on the local topographical and chemical heterogeneity.^4^

Not only the structure and dynamics of a protein change due to the presence of water, but also, the organization and dynamics of water molecules at the surface and interior of a protein are drastically modified depending on the local environment.^12,16–20^ Water is a deceptively simple molecule with many of its properties being highly context dependent. The physical behavior of water may vary depending on whether it is near a hydrophobic/hydrophilic molecule/surface, whether it is under confinement, and even different properties of water are perturbed to different extent depending on the nature of the perturbation.^21,22^

For water molecules bound to protein surface or buried inside protein cavities, the residence time may vary over a wide range depending on the nature of the interactions. Nuclear magnetic relaxation dispersion (NMRD) study by *Denisov et al*.^23^ revealed that the exchange time of water bound to protein may span over a broad range of time scales: 10 – 50*ps* for surface water and 0.1 – 1*ns* for those bound strongly to the protein. Even slower exchange time has been observed by Ernst *et al*^24^ in case of human interleukin-1*β* where they found water molecules trapped into hydrophobic cavities for longer than 1*ns*. An extensive theoretical study on the kinetics of water exchange in Cytochrome c by *Garcia et al*.^25^ depicts diverse time scales depending on the location of the binding site and the local interaction energies. A large number of computer simulation studies have elucidated the active role of bound water in and around proteins.^26–31^

In the present work, we explore in detail the unique structure, dynamics and energetics of the water molecules buried deep inside a photoactivated ion pump, namely KR2 rhodopsin from *Krokinobacter eikastus* in the resting (dark) state. Vision or photo-sensing in living organisms is possible due to the ability of rhodopsins to convert light into electrochemical signal in terms of translocation/pumping of protons/ions.^32,33^ Although the structure and function of a wide variety of rhodopsins have been known for a long time, they were mainly known to function either as inward chloride or outward proton pumps. Recently, the sodium ion (Na^+^) pumping activity of rhodopsin has been discovered for the first time in *Krokinobacter eikastus*.^34–36^

Similar to all microbial rhodopsins, KR2 is a transmembrane protein consisting of seven parallel *α*-helices that span across the membrane.^35–39^ A retinal chromophore is covalently attached to a lysine residue (LYS255) near the central part of the protein and undergoes cis-trans isomerisation upon activation through light and thus controls the ion transport across the cell membrane. KR2 contains a unique Asn-Asp-Gln (NDQ) motif which is used to pump Na^+^. ^34,40,41^ This is in contrast to bacteriorhodopsin (bR) which possesses highly conserved Asp-Thr-Asp (DTD) motif that carries out proton pumping.^42,43^ However, in the absence of Na^+^, KR2 acts as a proton pump. ^44^ Moreover, variants of KR2 have been designed to modulate electrical properties, substrate specificity with potential applications to optogenetics.^35,36,38,45,46^

Intuitively, one might speculate that the pattern of hydration along the ion pumping pathway should be quite different between the proton pumping bacteriorhodopsin (bR) versus Na^+^ pumping KR2. It has been widely accepted that translocation of protons is facilitated by a Grotthuss mechanism involving a chain of hydrogen bonded water molecules. On the other hand, for ion channels/pumps ion selectivity depends on the hydration dependent size of the ions. For example, hydrated Na^+^ ion has larger size than K^+^, whereas dehydrated Na^+^ has a smaller size. So it might be crucial for a Na^+^ to leave the water shell while passing through a selectivity filter. Interestingly, more recent crystal structures have suggested that KR2 may exist in a compact monomeric (in lower pH) and expanded pentameric (in slightly alkaline pH) states.^38,47^ The hydration level changes significantly between these two functional states. Since KR2 predominatly pumps proton at acidic pH, it is expected that the overall hydration structure could be similar to bacteriorhodopsin (bR), whereas at higher pH the hydration behavior would change facilitating the ion pumping. However, in this particular work we have used the monomeric form (PDB ID: 3X3B) to investigate the hydration pattern for computational efficiency.

Interestingly, all rhodopsins contain structurally conserved bound water molecules buried deep inside the protein channel. Hence, it has been often suggested that these bound water molecules may play a crucial role in the function of these biomolecules.^42,48-53^ The crystal structure of KR2 rhodopsin exhibits several such bound water molecules that motivates us to explore and characterise the nature of these buried hydration sites.^37,38,47^

Recent theoretical study on the localisation of water inside KR2 rhodopsin channel shows reorganisation of water molecules upon conformational transitions of the protein. ^54^ Another simulation study on Channelrhodopsin-2 (ChR2) revealed that the water acts as a lubricant in the ion transportation.^49^ In addition, the water at the specific sites may play a major role in the ion binding and transport processes.^42,55–57^ Therefore, understanding the molecular level structure and dynamics of water will help us to better understand the mechanism of Na^+^ pumping, and rhodopsins in general, which eventually could be used for the rational design of novel light-driven ion pumps and development of inhibitory optogenetic tools.^34,35,58,59^

### Computational details

We chose the PDB entry 3X3B as the initial structure for our simulation of KR2 protein in the dark state. ^35^ The water molecules that were resolved in the crystal structure and present along the ion conduction pathway were retained. The retinal was kept in the protonated state (positively charged). Standard protonation states were assigned to all other residues, i.e. all ASP/GLU were deprotonated (negative) and LYS/ARG were protonated (positive). Protonation states of histidines were optimized by the *pdb2gmx* utility of Gromacs.^60^ There were two histidine residues in the protein: HIS30 and HIS180. Both were in neutral states. They were protonated on the delta and epsilon nitrogens for HIS180 and HIS30, respectively. The protein was embedded and aligned in DMPC bilayer using the CHARMM-GUI webserver. ^61^ The major axis of the protein made an angle (~ 12°) with the *z*-axis, where the *z*-axis represents the perpendicular direction to the plane of lipid bilayer. The terminal residues were capped by acetyl (ACE) and N-methyl (NME) groups, respectively. The protein-membrane system was solvated by water (TIP3P water model) and neutralised by Cl^-^ ions. Additional Na^+^ and Cl^-^ ions were added to maintain 150 mM concentration of NaCl. The final simulated system comprised of 1 protein, 367 DMPC, 80 Na^+^ ions, 82 Cl^-^ ions and 26,776 water molecules. Periodic Boundary Condition (PBC) was employed in all directions. See Fig. S1 for a visual representation of the system.

The MD simulations were carried out using the Gromacs software suite (GROMACS v5.0.7).^60^ We used the CHARMM36m force field^62^ for all the components except the retinal chromophore. The force field for retinal was obtained from the earlier work by Hayashi *et al*.^63–67^ A cut-off radius of 1.2 nm was used for both the Lennard-Jones and coulombic parts of the potential. The long range electrostatic forces were taken care of by using the particle-mesh ewald (PME) method. ^68^ We carried out energy minimization using steepest descent algorithm. It was followed by a NVT equilibration run at T=310K for 1ns and NPT equilibration at T=310K and P=1bar for 10ns. The temperature and the pressure were maintained using velocity rescaling (v-rescale) thermostat^69^ and the Parrinnello-Rahman semi-isotropic barostat,^70^ respectively. The barostat was coupled separately along the *xy* plane and in the *z*-direction. All bonds with hydrogen atoms were constrained using LINCS algorithm which allowed us to use 2*fs* integration time step.

After the equilibration runs, the system was simulated at 310 *K* temperature and 1 bar pressure for 1.5*μs*. We carried out total three such independent simulations of 1.5*μs* each leading to a cumulative 4.5*μs* trajectory. For the analyses, we saved the frames at every 10*ps*, except for the computation of mean square displacement (MSD) and the rotational correlation function in which cases we deposited at every 1*ps*. For the computation of the residence time and the short time diffusion coefficient, we carried out additional 100 independent simulations (each 500*ns* long) starting with different initial configurations obtained from every 10*ns* time interval of the last 1*μs* of one of the three independent trajectories.

## Results and discussion

### Identification of hydration sites and extent of spatial localisation

In order to ascertain the stability of the protein structure during the course of the simulation, we have computed the RMSD of the protein backbone over a time period of 1.5 *μs* (Fig. 1A). The RMSD converges within hundreds of picoseconds to a value of 0.143 ± 0.11 *nm* and does not show any drift in the remaining simulation trajectory, thus, confirming the overall stability of the protein conformation.

**Figure 1:**
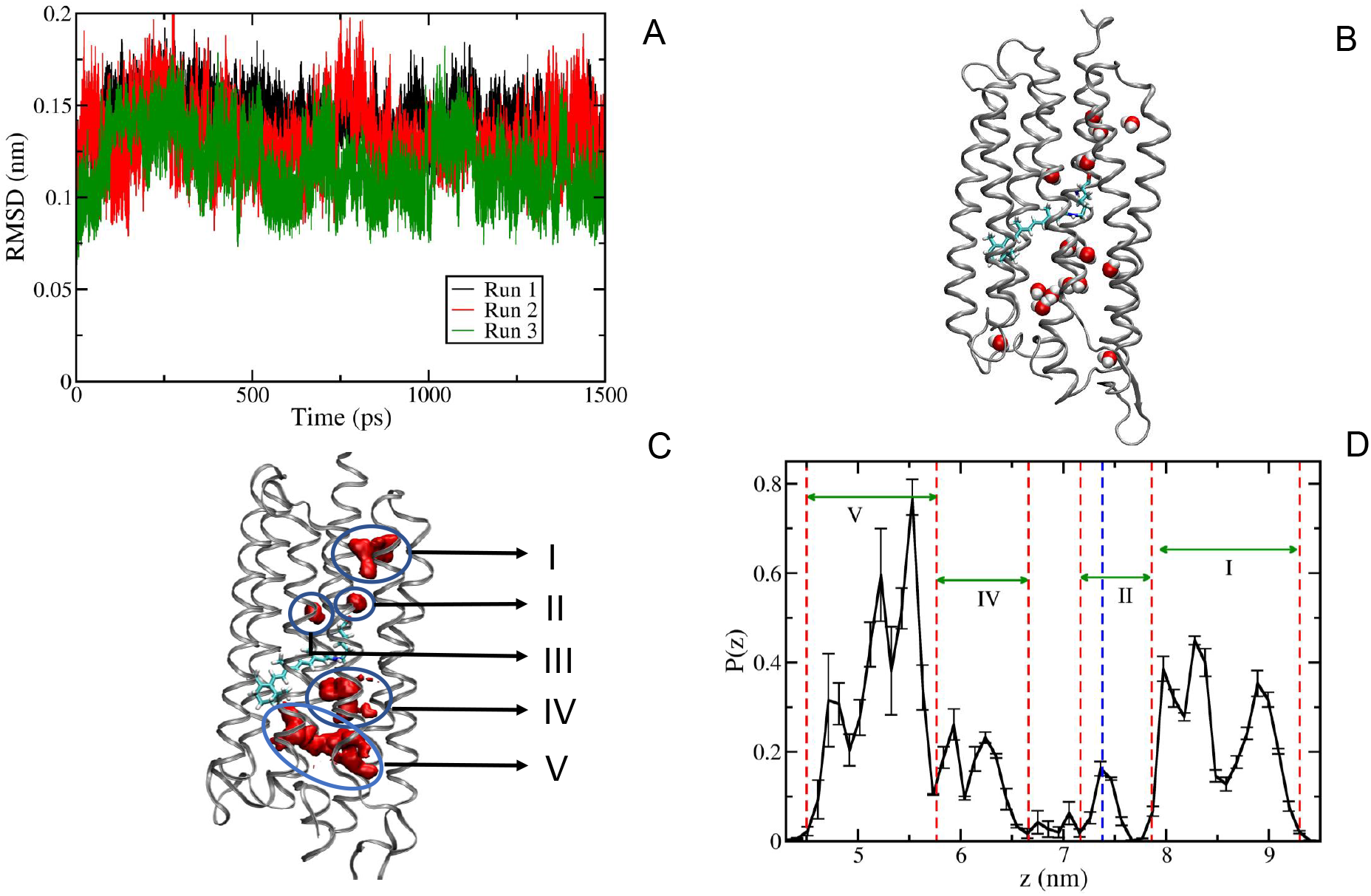
The stability of protein and spatial localization of water. (A) RMSD of the *C_α_* atoms of the protein as a function of time. The three data sets correspond to the three independent MD runs. (B) The crystal structure of KR2 rhodopsin in the resting state along with the retinal chromophore (cyan stick) and water molecules (red/white spheres) present inside the channel. (C) The spatial density map of water inside the channel is shown by red patches and distinct pockets/cavities are labeled from I to V. The cavities from I to III reside above the retinal chromophore towards the cytoplasmic side, whereas IV and V are below it towards the extra-cellular side. (D) The population density of water molecules along the *z*-direction (perpendicular to the membrane plane) confirms the existence of five distinct hydration sites along the channel. All the cavities except III are shown by regions enclosed by vertical red dashed lines. The position of the cavity III is indicated by the vertical blue dashed line. The error bars indicate the standard deviations of the average quantity obtained from three independent simulations.

Several crystal structures of KR2 contain multiple water molecules along the channel and they have been suggested to be functionally important.^35,38,47^ Earlier simulation study on KR2 rhodopsin has shown that even in the resting state water molecules may be continuously distributed along the channel forming two clusters separated at the vicinity of the retinal. ^54^ In their work, discrete positions of water molecules at different time frames were superimposed to visually identify the clusters. Instead of that, in order to generate a more refined statistical measure of the water distribution, we have computed the spatial density of water along the channel. The spatial density has been computed by averaging the local densities (in three dimensional grids) along the channel over the 1.5*μs* long MD trajectory. The density of water clearly shows five distinct pockets/cavities (I-V) along the channel (Fig. 1C).

The probability distribution of water molecules along the *z*-axis (perpendicular to the membrane plane) further demonstrates the localisation of water at five specific sites of the channel (Fig. 1D). This is consistent with the localisation of water molecules in the crystal structure^35^ (Fig. 1B) and recent experimental observations.^38,47^ Based on the location of the cavities, we have labeled them from I to V from the intracellular (top) to the extracellular (bottom) side (Fig. 1C). While the cavities I, IV and V appear to be broad, the cavities II and III are significantly small in size. We observed that the cavity III contains only one water molecule throughout the 1.5*μs* simulation time, whereas the number of water inside the cavity II fluctuates between 1-2. To verify the significance of the result in Fig. 1D, we have carried out two additional MD runs of length 1.5*μs* each. The results are shown in Fig. S2 in the supplementary material. Consistent with the earlier studies, we have observed a separation between the water clusters by ~ 6Å at the vicinity of the retinal (note the gap between the cavities II and IV in Fig.1D)).^49,54^

### Hydration level of the cavities

The average number of water molecules within the first hydration shell (coordination number) of a water in bulk is 4.5 (Fig. 2A). ^25^ A water in the surface of a biomolecule is expected to have reduced number of neighbouring water due to the limited accessibility imposed by the biomolecule. This number should vary significantly for water molecules inside a cavity depending on the size and nature of the cavity, i.e. the amount of water present in it. In KR2 rhodopsin, all the cavities identified in our study have severely reduced coordination number suggesting reduced hydration level inside the cavity. For all the cavities except II and III, we find that the average number of water molecules within the first hydration shell is between 1-3. For cavities II and III this number is ~0. Thus, the overall hydration level of water inside rhodopsin channel is significantly compromised due to the confinement.

**Figure 2:**
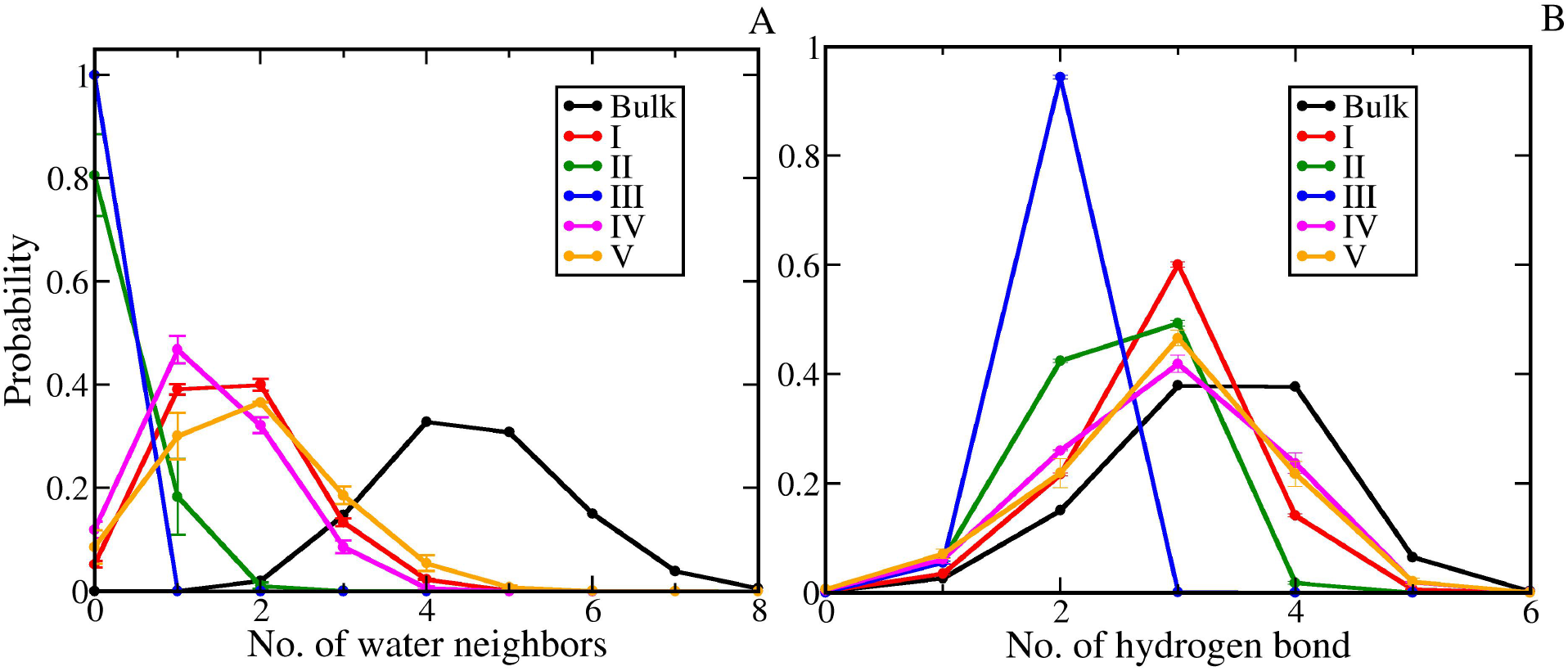
Distribution of the number of neighboring water molecules and the number of hydrogen bonds (both with water and protein) for the water molecules in different cavities inside the channel in comparison to the bulk water. (A) The probability distributions of the number of neighboring water molecules in the first hydration shell (radius 3Å) of the trapped water molecules show a drastic depletion in number compared to that of the bulk water indicating reduced hydration. The water molecule in the cavity III does not have any neighboring water. (B) The probability distributions of the number of hydrogen bonds for the trapped water molecules are similar to that of the bulk water except the one in the cavity III which shows significantly less number of hydrogen bonds. The error bars indicate the standard deviations of the average quantity obtained from three independent simulations.

Although the hydration level in the cavity is compromised, the number of hydrogen bonding partners (including both water and protein residues) remains comparable to that of bulk water for all the cavities except cavity III (Fig. 2B). The average number of hydrogen bonds in these cavities is ~ 3 compared to ~ 3.5 in case of bulk water. Hence, the energetic penalty due to desolvation can be to some extent compensated by making additional hydrogen bonding interactions with the surrounding amino-acid residues. But for cavity III, the average number of hydrogen bond is ~ 2. Thus, for cavity III, not only hydration level, but also the number of hydrogen bonding partners is significantly lower. We have carried out the same analyses for two additional independent MD trajectories of length 1.5*μs* each. The results are shown in Figs. S3-S4 in the supplementary material.

### Heterogeneity in the water-protein interaction energy within cavity

The interaction energy of the water molecules in cavities I-III with the protein shows normal distribution indicating homogeneous environment in energy space (Fig. 3A). On the contrary, the water molecules in cavities IV and V display a broad distribution in energy suggesting a heterogeneous environment. The probability distribution of the water molecules in the cavity V shows a broad distribution over a wide range of protein-water interaction energy. Interestingly, in case of cavity IV, the distribution shows two distinct peaks. These peaks are observed to be associated with different environments as described below. In addition, the water-water and water-protein interaction energies for the trapped water molecules are found to be anti-correlated to each other (Fig. 3B). It is to note that we have carried out additional two independent MD simulations of length 1.5*μs* each to verify that the results are statistically significant (see Fig. S5).

**Figure 3:**
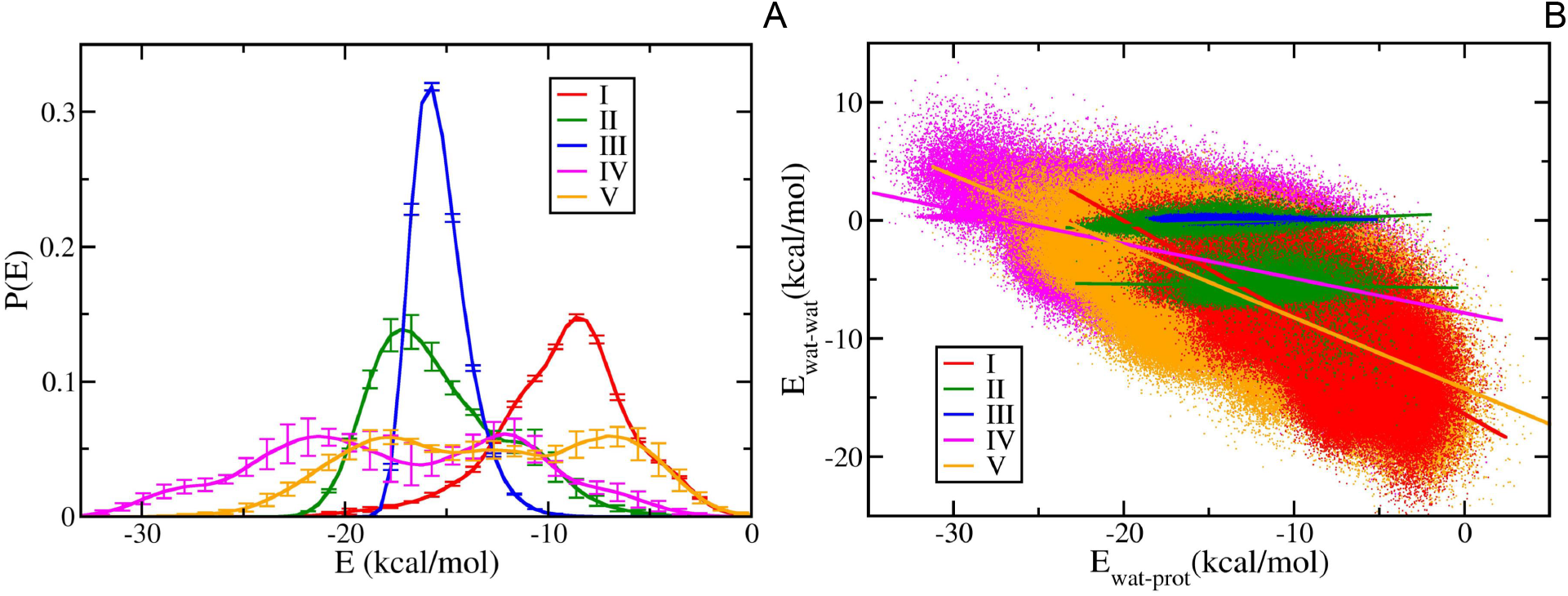
Interaction energy of trapped water molecules with the environment. (A) The water molecules inside the cavities situated above the retinal chromophore (cavities I-III) show a Gaussian distribution in the water-protein interaction energy suggesting homogeneous environments in terms of interaction with protein. On the contrary, the water molecules trapped in the cavities situated below the chromophore (cavities IV and V) show a broad non-Gaussian distribution indicating heterogeneous environment. Interestingly, the water in the cavity IV shows two distinct peaks. The error bars indicate the standard deviations of the average quantity obtained from three independent simulations. (B) The water-water (E_wat-wat_) and water-protein (E_wat-prot_) interaction energies are anti-correlated to each other for the trapped water molecules except those trapped in the cavities II and III. The lack of such anti-correlation in these cavities is due to low hydration level.

### Protein-water interaction pattern in the cavities

The local environments of the water molecules trapped in each of the cavities are depicted in Fig. 4. The cavity I contains a cluster of water molecules involved in a hydrogen bonded network (Fig. 4). A similar structure has been observed earlier in bacteriorhodopsin. ^71^ The cavity I is surrounded by key residues T49, I50, S60, N61, S64 and G259 (see Table 1). It is in the cytoplasmic side and the water in this cavity is easily exchanged with the cytoplasmic water. The location of this cavity is consistent with the ion-uptake cavity (IUC) identified in more recent crystal structures.^38,47^ This particular cavity has also been referred as the “ion-selectivity filter”, since mutations in N61 and G263 allowed KR2 to pump larger ions like K^+^.^72^

**Figure 4:**
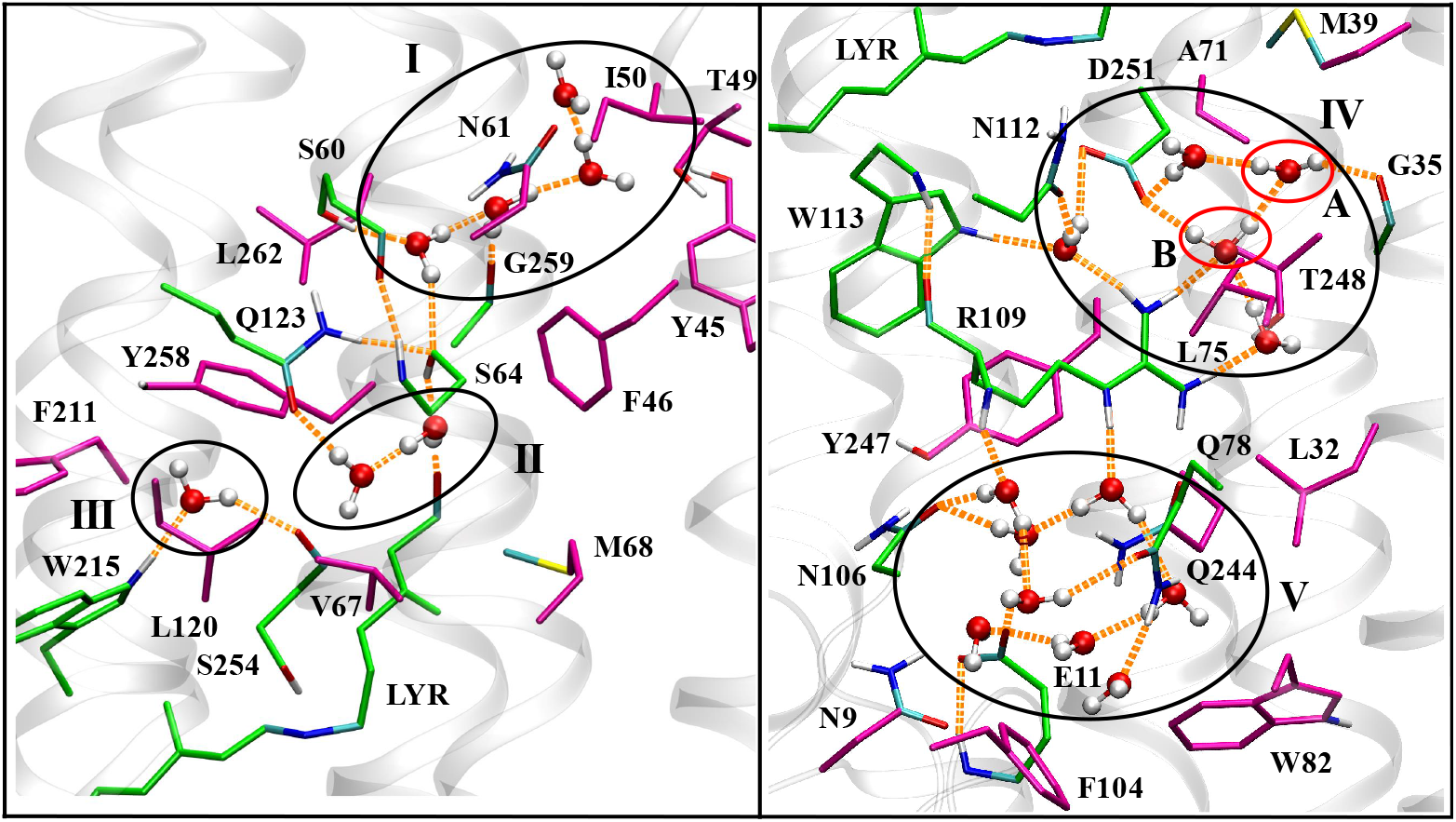
Local environments of the water molecules trapped in each of the cavities. Left panel: The water molecules trapped in the top part (above retinal) of rhodopsin (cavities I, II and III). The water molecules in cavity I show a cluster of hydrogen bonded water molecules. The water molecules in cavity II are surrounded by S64, Q123 and LYR (Lysine attached to retinal). The water trapped in the cavity III is surrounded by Y258, F211, W215 and S254. It forms strong H-bonds with W215 and S254. Right panel: The water molecules those are trapped in the bottom part of rhodopsin (cavities IV and V). The cavity IV is large in size and it accommodates multiple water molecules. This cavity is heterogeneous in terms of interaction between water and protein (see Fig.3A). We have identified two local environments (labeled as A and B) with high residence probability of water. The cavity V resides at the exit part of the channel and it is connected to the bulk water. The key residues defining each cavity are listed in Table 1.

**Table 1:** List of amino-acid residues surrounding the cavities

| Cavity | Surrounding residues |
| --- | --- |
| I | Y45, T49, I50, S60, N61, S64, G259, L262 |
| II | L43, F46, S64, V67, Q123, LYR255, Y258 |
| III | L120, F211, W215, S254, Y258 |
| IV | G35, M39, A71, L75, R109, N112,<br>W113, Y247, T248, D251, LYR255 |
| V | N9, E11, N12, T28, L32, Q78, N106, R109, Q157,<br>E160, Q244, Y247, W82, F104 |

The cavity II is constituted by F46, S64, Q123, LYR255 and Y258 and is situated below the cavity I (Fig. 4). The water in this cavity is stabilized by formation of hydrogen bonds with S64, Q123 and LYR255. Although the Fig. 4 shows two water molecules in this cavity, the number of water in the cavity II fluctuates between 1-2. The presence of an extended hydrogen bonded network involving protein residues (including Q123) and water in this region has been investigated extensively by Tomida *et al* using difference FTIR spectra.^72^ These studies have established that certain mutations in this region, e.g. Q123A, Q123V, and S64A disrupt the pumping action due to perturbation of this important hydrogen bonded network.

The cavity III is surrounded by F211, W215, Y258 and S254. The water in this cavity forms two hydrogen bonds with amino acid residues S254 and W215. Cavity III contains only one water molecule, which is highly conserved across independent crystal structures of KR2. Interestingly, water in this site has exceptionally high residence time and remains in the cavity for the entire duration of 1.5*μs* MD trajectory as will be demonstrated later (Fig. 5). However, the functional importance of this site/water molecule remains unknown. We speculate that this water might be responsible for holding the residues S254 and W215 in the right orientation by acting as a bridge. This aspect will be further highlighted later in terms of the highly anisotropic rotational dynamics of this particular water in cavity III (Fig. 8).

**Figure 5:**
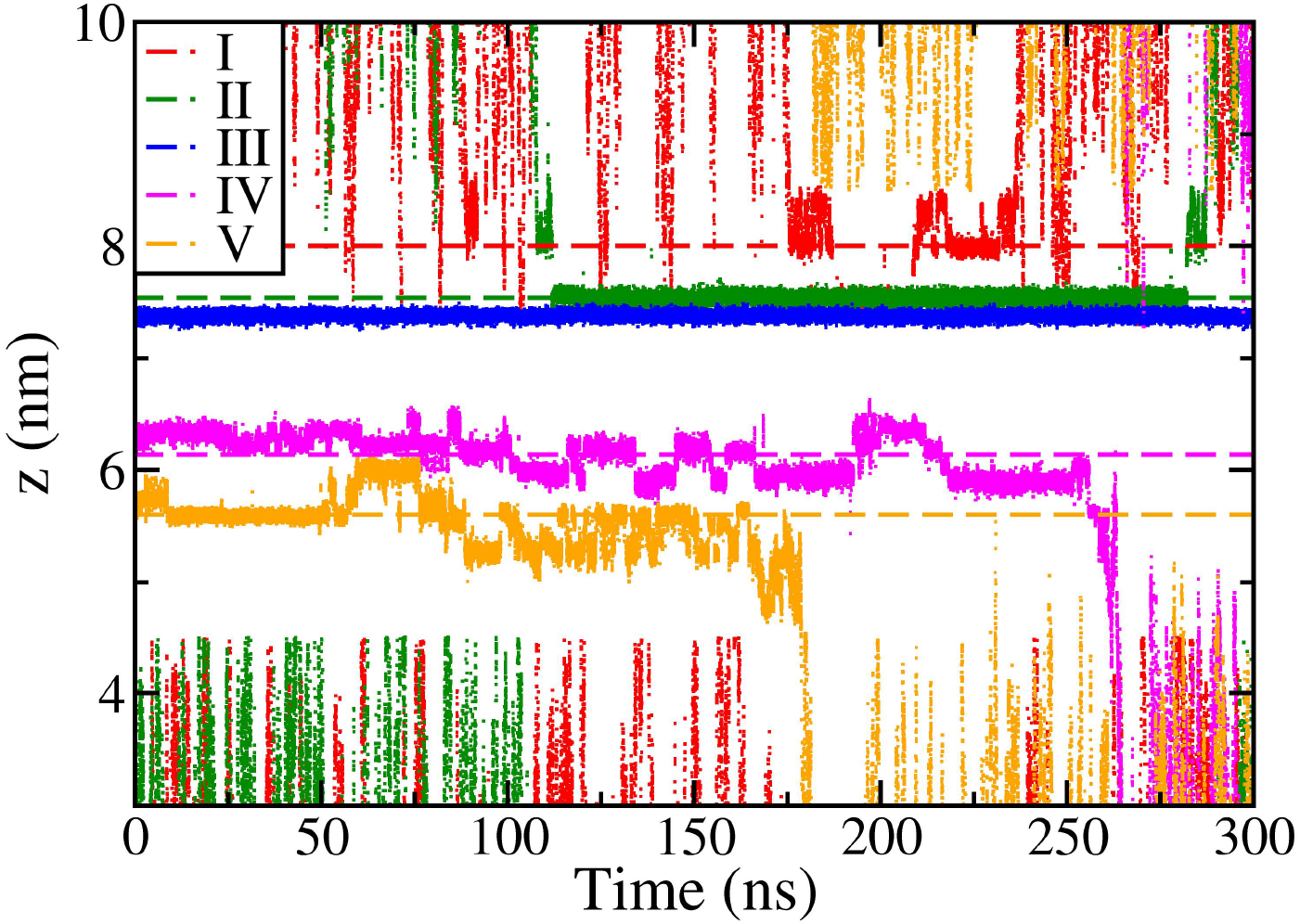
The z-components of positions of few representative water molecules that are trapped inside the cavities. The trapping events are evident from the reduced fluctuations in the z-values. The dashed lines represent average locations of respective cavities. The significantly long residence times in a few cavities are prominently visible.

The cavity IV is much bigger in size and therefore contains multiple water molecules. Consistent with the energetic heterogeneity as shown in Fig. 3A, there are two distinct local environments (Fig. 4, right panel). These two types of water molecules have following hydrogen bond partners: (i) water in region A forms hydrogen bond to the backbone of G35 and (ii) water in B shares hydrogen bonds with two neighboring amino acid residues D251 and R109. Thus, the number of amino-acid partners increases from A to B in accordance with the interaction energy distribution as shown in Fig. 3A. In addition, there are few other water molecules present in the cavity with varying number of water-water hydrogen bonds. This observation is consistent with the IR spectroscopic analysis of KR2 in the dark state. ^73^

The cavity V resides at the exit of extracellular side. This is more like a channel containing several water molecules having diverse local environments. Some of the residues that form this cavity are listed in Table 1. This particular cavity coincides with the “ion release cavity” identified by crystallographic studies.^38,47^

### Residence time distribution in the cavities

As observed earlier, the water on a biological surface or inside cavity can stay for long enough time ranging from a few picoseconds to microseconds. Not only they reside inside the channel, the water molecules take fairly long time (~ 100 *ns*) to enter into the channel (see Fig. S6). In order to obtain the residence time of trapped water in different cavities, we have followed the *z*-coordinate of the O-atoms of the water molecules over time (Fig.5), where *Z*-axis is perpendicular to the membrane plane. The scattered dots above and below the membrane bilayer signify the diffusive motion of the water molecules in the bulk phase. Occasionally they bind to the protein/membrane surface and get trapped into cavities (depicted by regions with *z* values fluctuating around a constant value with narrow width). The positions of the identified cavities are marked by horizontal dashed lines of same color.

We find a diverse range of residence time of trapped water molecules depending on the identity of cavity. The water molecules in cavity I dwell for relatively short period of time ranging from 1 to 10 ns. On the other hand, those in the cavities II, IV and V remain trapped for longer period of time (tens to hundreds of nanoseconds). Surprisingly, the water in cavity III remains trapped during the entire simulation time scale of 1.5 *μs*! Thus, this cavity is very unique in nature. It does not contain any neighboring water molecule; remains in an isolated trapped environment and never gets exchanged with water in the other cavities. A similar experimental observation was reported earlier in a different system.^53^ We speculate that this particular highly conserved water molecule may have a significant functional role during the photocycle that is yet to be elucidated.

Table 2 reports the average residence time of water molecules in each cavity. The averages were obtained over all the water molecules that entered the respective cavities during the course of the simulation. As the table suggests, the cavities near the membrane boundaries (I, II and V) are easily accessible to the bulk water and therefore they show relatively short mean residence time ranging from few nanoseconds to tens of nanoseconds. On the other hand, the water molecules in the relatively deeply buried cavity (IV) stay trapped for hundreds of nanoseconds. The water inside the cavity III never escapes out of the cavity within our simulation timescale.

**Table 2:** List of logarithmic average residence time and diffusion constant in different cavities including in bulk. The residence time (*τ*) is in *ns* and the diffusion coefficient (*D*) is in *cm*^2^/*s*.

| Cavity | $\log(\tau)$ | $\log(D)$ |
| --- | --- | --- |
| Bulk | - | -3.5 |
| I | $0.51 \pm 0.09$ | $-5.8 \pm 0.07$ |
| II | $0.90 \pm 0.09$ | $-7.0 \pm 0.16$ |
| III | $> 3.2$ | $-\infty$ |
| IV | $1.8 \pm 0.15$ | $-6.6 \pm 0.15$ |
| V | $1.15 \pm 0.32$ | $-5.9 \pm 0.16$ |

Fig. 6A shows the probability distribution of residence time of water in each cavity in logarithmic scale. The escape events are random as justified by the log-normal nature of the distributions, which is ubiquitous in nature. ^74^ This is to note that an earlier study by Garcia *et al* in Cytochrome c reports a power-law decay of probability with increasing residence time with the exponent value ranging from 0.4 to 1.8.^75^

**Figure 6:**
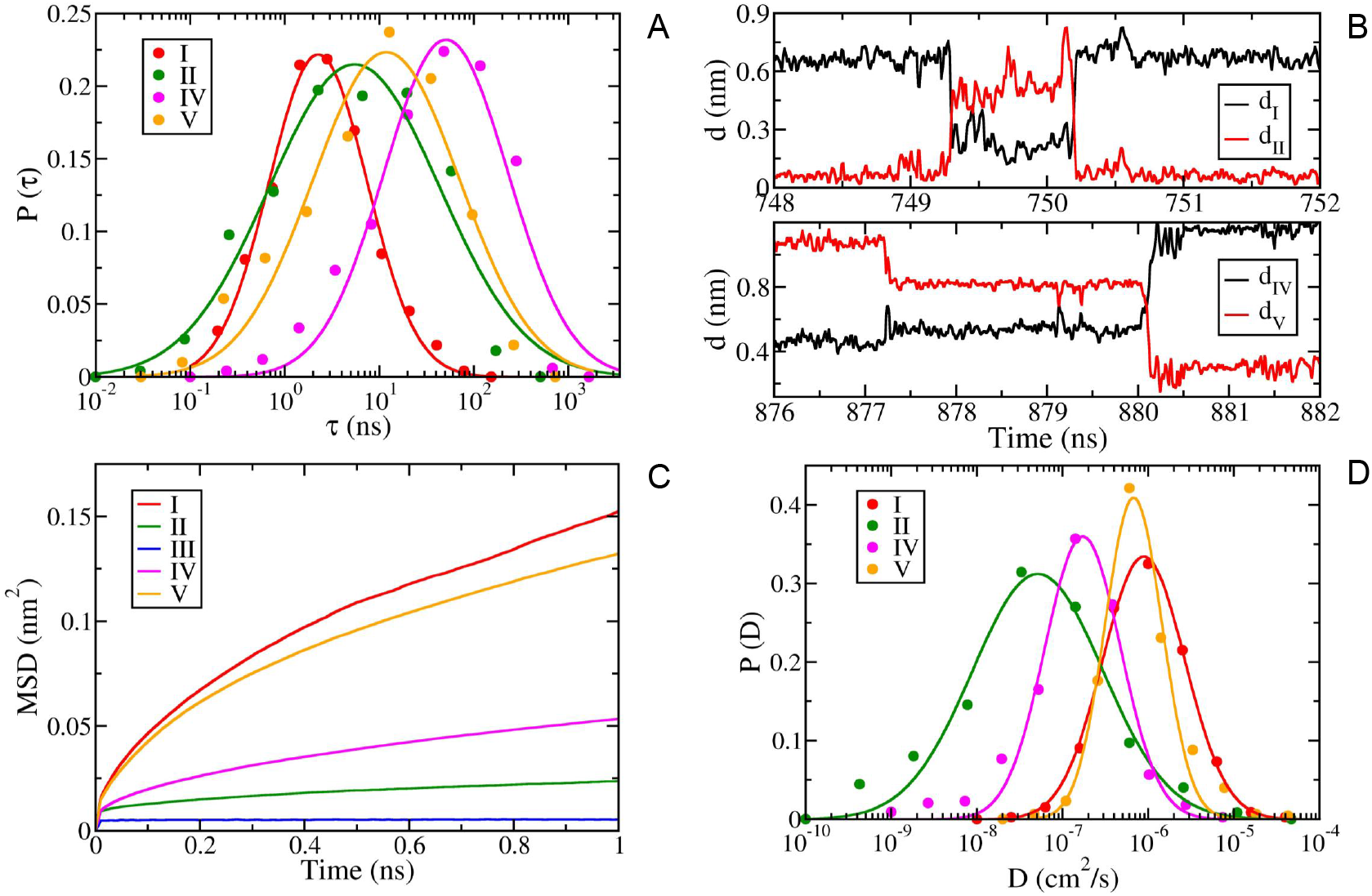
(A) Probability distribution of residence time of water molecules inside different cavities along the channel of KR2. It shows a broad range of residence time starting from tens of picoseconds to micro-second, consistent with the broad distribution of energy associated to the water molecules in different cavities as shown in Fig.3A. (B) The transitions of water molecules in between cavities happen through jump-like motion (activated process). The top panel shows the time evolution of distances of a representative water molecule from cavity I (black line, *d_I_*) and II (red line, *d_II_*). The bottom panel shows the distances of another representative water molecule from cavity IV (black line, *d_IV_*) and V (red line, *d_V_*). (C) Mean square displacement (MSD) of water molecules inside the cavities shows a diffusive motion at short times (< 1*ns*) as indicated by linear growth of MSD with time. (D) The probability distribution of the short-time diffusion constants of water molecules within the cavities: Diffusion coefficients were computed from the slope of the linear regime of time dependent MSD curves.

### Bottlenecks separating the cavities

The long residence times of each of the cavities provide an evidence of presence of bottlenecks separating the cavities. We find that the residues Q123 and S64 act as gates in between cavities I and II consistent with earlier crystallographic studies.^38,47^ The water molecules in cavity I are well hydrated and form a hydrogen bonded network. In order to escape and enter into cavity II, they need to break the hydrogen bonds in cavity I and then cross the bottleneck created by Q123 and S64. These two residues form hydrogen bonds with water molecules that try to escape cavity I and enter cavity II, thus create a gateway that creates a barrier to such transitions. This observation is particularly relevant in the view of the several mutations in Q123 that disrupt the pumping mechanism of KR2.

The residues S254, Y258 and W215 create an isolated environment for the water in cavity III. Furthermore, the strong hydrogen bonds formed by S254 and W215 with the water within such confined environment completely isolates the water. Therefore, water from cavity II cannot enter cavity III and vice versa. There is no exchange of water in between cavities II and IV as they are well separated by retinal (Fig. 1C). The water in cavity II cannot reach retinal due to the presence of L120 and V67 that act as hydrophobic gate-keepers as observed in the previous study by Kaila and coworkers. ^54^ The cavities IV and V are separated by mainly R109 that controls the exchange of water in between them. Thus, overall the water exchange between cavities are controlled by few key amino-acid residues along the ion channel pathway.

The exchange of water between the cavities occur through a jump-like motion (hopping) due to the presence of a free energy barrier created by the bottlenecks. A few representative trajectories of such transitions are shown in Fig. 6B for transitions between cavities I and II (top panel) and cavities IV and V (bottom panel).

### Heterogeneity in translational motion and diffusivity

The dynamics of bound water molecules strongly depends on their local environments. As we have observed the water molecules inside the rhodopsin channel are quite different from that in the bulk in terms of their local environment, their dynamics is expected to be different as well. To gain an insight about the translational motion, we have computed the mean square displacement (MSD) as a function of time for water in each cavity (Fig. 6C). The water molecules in cavities I, IV and V show diffusive behavior within nanosecond as indicated by the linear increase of MSD with time. However, their diffusion constants are markedly lower compared to that of bulk water (Table 2). In contrast, the water molecules in cavities II and III show signature of significantly trapped motion indicated by rapid saturation of MSD. The MSD values at the saturation provide an approximate measure of the respective cavity size. The cavity sizes for II and III are estimated to be ~0.15 nm and ~0.07 nm, respectively, that are smaller than the size of a water molecule (~0.35 nm) indicating severe motional restriction. We must note that the larger cavities (I, IV and V) are expected to show the signatures of trapped diffusion as well had we computed MSD to longer times.

The environment of bulk water is homogeneous and identical for every water molecule. On the contrary, each time a water enters a cavity it faces a different environment for the finite duration of time it stays inside the cavity depending on the species it is interacting with. This leads to a significant heterogeneity in the diffusive motion at a molecular level even within the same cavity. As shown in Fig. 6D, the diffusion constant computed at single molecule level for each cavity shows a broad distribution. The log-normal distribution of diffusion constant is Gaussian in nature and shows a variation in three orders of magnitude (~ 10^−9^ to ~ 10^−6^). This is to note that the water in the cavity III is almost frozen and therefore we have not computed the corresponding distribution.

### Heterogeneity in rotational dynamics

Confinement in cavities is expected to significantly hinder the rotational relaxation of the trapped water molecules. Water on bio-molecular surfaces with slow translational motion is generally associated with slow rotational dynamics as well. Consistent with this fact, the trapped water molecules in all the cavities show slower rotational dynamics compared to bulk water as reflected in the slow decay of the first rank rotational time correlation function (Fig.7) computed using the following equation:

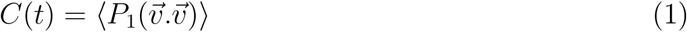

where, *P*_1_(*x*) = *x* is the first rank Legendre polynomial and 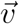 is the unit vector perpendicular to the water plane. We have fitted the data with the following tri-exponential form to identify the distinct time scales and their contributions to the overall rotational dynamics.

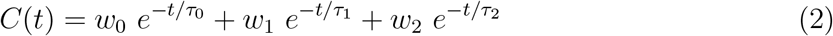

where, *w*_0_, *w*_1_ and *w*_2_ are the weight factors and *τ*_0_, *τ*_1_ and *τ*_2_ are corresponding time constants. The best fit parameters are reported in Table 3.

**Figure 7:**
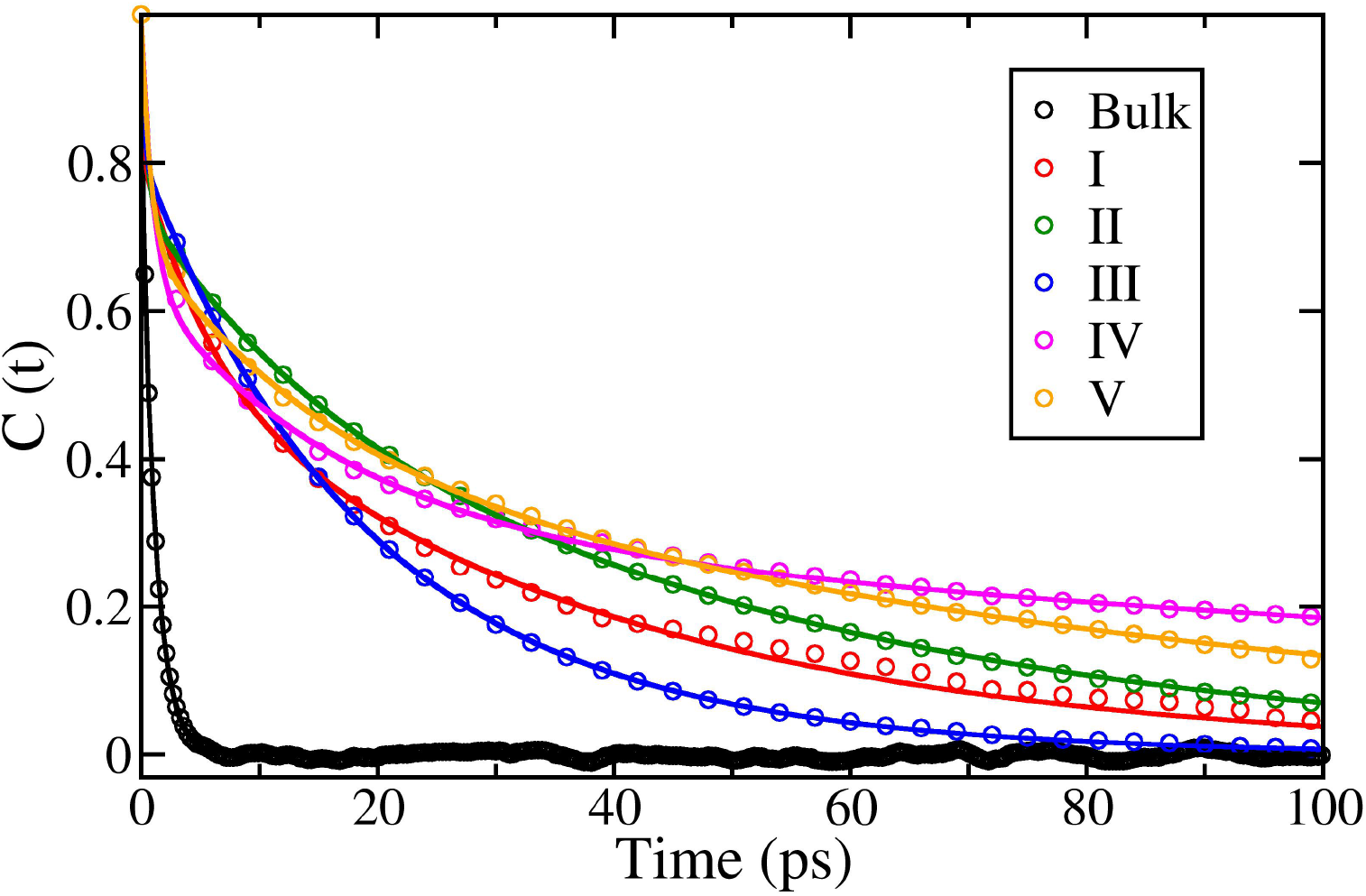
Rotational dynamics of water molecules in various cavities in comparison to bulk. Rotational time correlation functions of trapped water molecules inside the cavities show that the rotational relaxations in all the cavities are much slower (tens of picoseconds) compared to that of bulk water (~ 1 ps). Note that the water in cavity III shows rotational relaxation time of similar range in comparison to the water molecules in the other cavities. This clearly indicates that the environment in cavity III does not impose any additional constraints on rotational motion despite of its frozen translational motion.

**Table 3:**
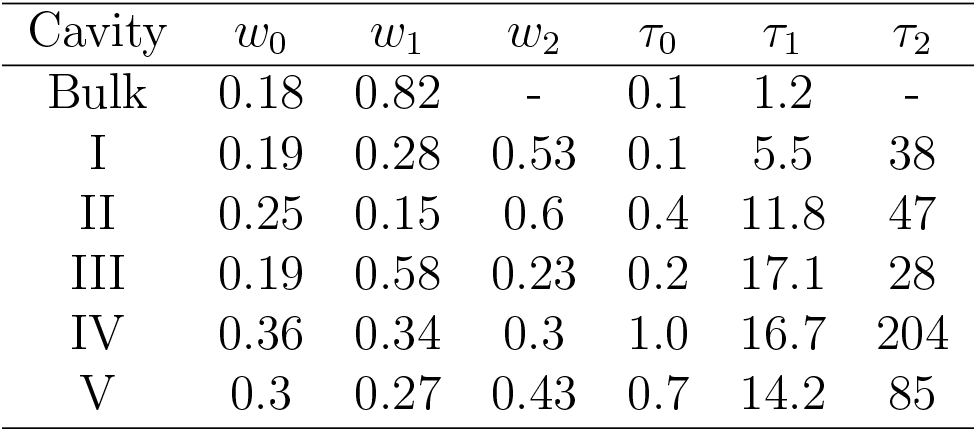
Rotational time scales (time constants are in picosecond)

| Cavity | $w_0$ | $w_1$ | $w_2$ | $\tau_0$ | $\tau_1$ | $\tau_2$ |
| --- | --- | --- | --- | --- | --- | --- |
| Bulk | 0.18 | 0.82 | - | 0.1 | 1.2 | - |
| I | 0.19 | 0.28 | 0.53 | 0.1 | 5.5 | 38 |
| II | 0.25 | 0.15 | 0.6 | 0.4 | 11.8 | 47 |
| III | 0.19 | 0.58 | 0.23 | 0.2 | 17.1 | 28 |
| IV | 0.36 | 0.34 | 0.3 | 1.0 | 16.7 | 204 |
| V | 0.3 | 0.27 | 0.43 | 0.7 | 14.2 | 85 |

The bulk water shows only two time scales in which the time constant 1.2 ps is the major contributor. In contrast, the trapped water shows three distinct time scales: (i) 0.1-1 ps, (ii) ~ 1-20 ps and (iii) hundreds of picoseconds. The largest time scale is expected to be associated to the confinement and the interaction of water with surrounding amino acid residues which are absent in the case of bulk water. Interestingly, the slowest time scales for cavities IV and V (below the retinal) are much larger than that of the cavities I-III (above the retinal) which suggests that the water at the upper part of retinal in the resting state are rotationally more flexible than the bottom counterpart. Thus, they are expected to have different dielectric response to the moving ions. The observation of faster rotational time scales for cavities II and III compared to that of IV and V is somewhat surprising, since we have seen earlier that the water molecules in cavities II and III are translationally trapped. Thus, some specific interactions in these cavities lead to decoupling in the translational and rotational dynamics of the water molecules.

We find a rather unique rotational dynamics for the water frozen in cavity III. The first rank rotational correlation function (Eq.1) of the vector perpendicular to water plane decays to zero in ~ 100*ps*. We computed the second rank rotational correlation function using following equation:

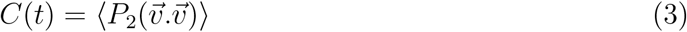

where, 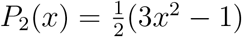 is the second rank Legendre polynomial and 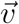 is the unit vector of concern. Interestingly, it does not fully decay to zero in ~ 100*ps*. Rather it saturates to ~0.6 at long time (Fig.8A). This suggests that the rotation in this cavity is not a free motion and could be highly anisotropic in nature. In order to investigate the reason behind it, we computed the time dependent *O_w_* — *H_w_* — *O*_*S*254_ angle and it reveals a frequent large rotational jump motion (Fig. 8B). The probability distribution of this angle appears to be bimodal in nature (Fig. 8C). The local configuration corresponding to the most probable angle is shown in Fig. 8D. The flipping of the water molecule exchanges the H-bonding atom with S254, while keeping the H-bond with W215 intact. Thus, the rotational motion of this water in the cavity III is highly unusual and has very specific coordinated flipping rotational motion.

**Figure 8:**
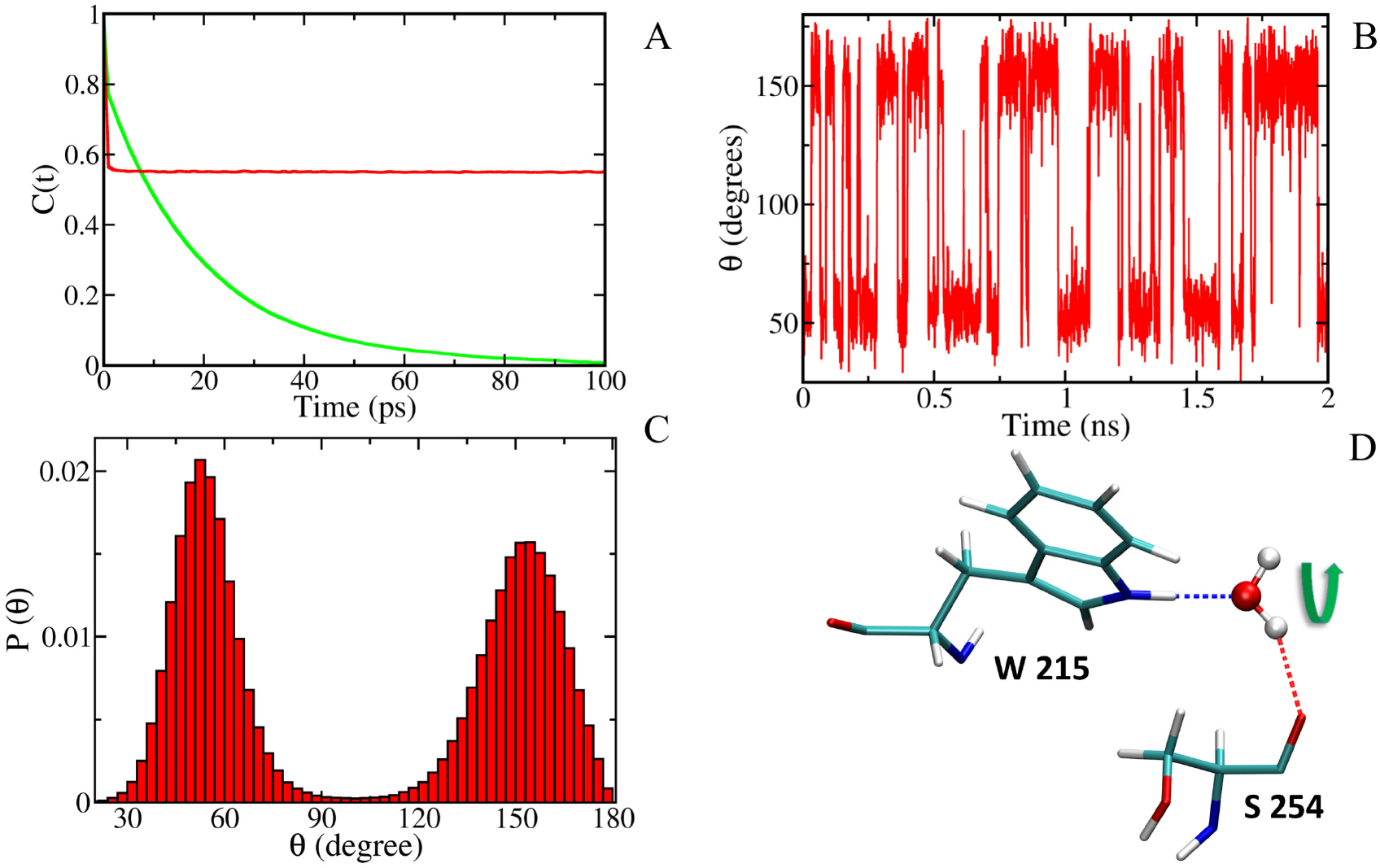
The anisotropic rotational dynamics of water molecule trapped in the cavity III. (A) Rotational time correlation functions of trapped water inside the cavity show that although the *P*_1_ (first rank Legendre polynomial; green line) decays to 0, the *P*_2_ (second rank Legendre polynomial; red line) does not decay to 0 rather it saturates at ~ 0.6 suggesting that the rotation is not completely free motion. (B) The time dependence of *O_w_* — *H_w_* — *O*_*S*254_ angle shows frequent large fluctuations. (C) A bimodal distribution of the probability of the *O_w_* — *H_w_* — *O*_*S*254_ angle is observed indicating presence of a bistable potential. (D) The local environment of the water molecule shows formation of two H-bonds with S254 and W215 residues. The water frequently rotates as depicted by the arrow by transiently breaking the H-bonds and then forming it again.

### Rotational anisotropy

Although the translational motion of a water in the bulk is isotropic in nature, the rotational motion as shown in Fig. S7 is observed to be slightly anisotropic (*I_A_*=2, where *I_A_* is the ratio of the largest and the smallest rotational timescales among the three rotational correlation functions computed for three orthogonal vectors: along dipole moment of water, along H-H direction and the vector orthogonal to the water plane). This is consistent with the earlier study by Ropp *et al*.^76^ Since each of the three rotational correlation functions is multi-exponential in nature, we have obtained the effective time scale for each of them using following equation:

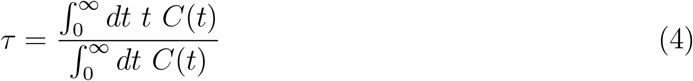

The rotational motion of water on biological surface is slowed down compared to that in the bulk.^6,77–79^ We find a similar slowing down for the water molecules inside cavity. In addition, we observed that the rotational motion inside protein cavity is highly anisotropic with the rotation of the dipole moment vector being the slowest among all the three orthogonal vectors (Fig. S7). The anisotropy indices of water molecules in the cavities I and V are 11 and 14, respectively (see Table 4). The rotational dynamics is more anisotropic as one moves towards the core of the protein. The water molecules in the core cavities II and IV show anisotropy indices 40 and 24, respectively. On the other hand, the water in the cavity III is extremely anisotropic in nature as discussed in previous section with observed anisotropy index ~ 10^12^!

**Table 4:** Rotational time scales of three mutually orthogonal vectors and rotational anisotropicity. Time constants are in picosecond.

| Cavity | $\tau_{HH}$ | $\tau_\mu$ | $\tau_\perp$ | $I_A$ |
| --- | --- | --- | --- | --- |
| Bulk | 15.03 | 21.3 | 10.4 | 2 |
| I | 42.83 | 544.93 | 37.74 | 14 |
| II | 52.54 | 1715.68 | 42.23 | 40 |
| III | 15.41 | $5.17 \times 10^{13}$ | 15.41 | $3.35 \times 10^{12}$ |
| IV | 64.44 | 1266.04 | 53.45 | 24 |
| V | 80.59 | 902.82 | 90.72 | 11 |

## Conclusion

We have carried out extensive all-atom molecular dynamics simulations to explore the structure and dynamics of water inside rhodopsin (KR2) channel in the dark state. Our study reveals five specific regions where the water molecules are trapped for significantly long time. These regions/cavities are spatially disjoint indicating the presence of protein side-chains acting as gateways controlling the movement of water between these cavities.

The cytoplasmic side (above the retinal chromophore) possesses three distinct cavities (I, II and III) whereas the extracellular side contains two cavities (IV and V). Cavity I is well connected to the cytoplasm and there is rapid exchange of water molecules. In this “ion uptake cavity” multiple water molecules are accommodated and they form a string like extended hydrogen bond network in agreement with earlier FTIR studies. The cavity II, located below the cavity I and well separated from it, can accommodate up to two water molecules. The sodium ion must reduce its hydration shell in order to jump from cavity I to II through the ion selectivity filter. The protein-water hydrogen bonded network around the Schiff base is crucial for the ion pumping activity. The cavity III carries strictly one water molecule that has unusually high residence time (> 1.5*μs*) and high rotational anisotropy. It holds two key protein residues (W215 and S254) together through strong hydrogen bonds. The cavities IV and V accommodate several water molecules and they constitute the exit path of the pumped ion.

The local structure and dynamics of these water molecules demonstrate significant heterogeneity. The water-protein interaction energy is highly heterogeneous indicated by the non-Gaussian nature of the energy distribution. The cavity IV shows appearance of two distinct peaks in the interaction energy distribution. The observed residence time of trapped water varies from few nanoseconds to hundreds of nanoseconds depending on the identity of the cavity. Interestingly, the water inside cavity III never escapes in our 1.5*μS* simulation time, which is extremely unusual!

The translational motion of the trapped water is diffusive in nature within 1 ns but is markedly slowed down compared to the bulk as expected. Water molecules in the cavities II and III show signature of trapped diffusion with the water in cavity III being almost frozen in terms of translational diffusion. Although the water molecules diffuse within the cavities, the exchange of water in between cavities is rare and it occurs through hopping motion controlled by the protein sidechains acting as gateways. Thus, the movement of ion is expect to occur through similar jumps between these cavities, since these are likely to serve as intermediate ion binding sites.

The rotational dynamics of the buried water molecules show multi-exponential decay with wide range of timescales. The slowest component of the decay is of the order of hundreds of picoseconds as compared to ~1 picosecond in case of bulk water. Thus, the rotational dynamics of trapped water is slowed down by a factor of ~100. Water in the cavity III depicts a unique flipping motion between two distinct orientations. We find the rotation of the trapped water molecules to be highly anisotropic in nature.

The results presented in this study lay the groundwork for understanding the intimate role that hydration might play in the sodium-pumping mechanism of KR2 rhodopsin. The hydration sites are likely to serve as the intermediate ion binding sites and control the energetics and kinetics of the ion pumping. It would be interesting to explore how the nature of these hydration sites and their connectivity pattern vary in subsequent stages of the photocycle (after photoactivation). While this work focuses exclusively on the water structure and dynamics in the resting state of KR2 rhodopsin, future work will further explore the coupling between the water and protein dynamics, and their effects on the various stages of the overall photocycle.

## Supplementary Material

See supplementary material for additional figures and analysis data supporting the conclusions drawn in this work.

## Acknowledgement

SC thanks SERB/DST, India for funding (ECR/2018/002903). Authors thank SNBNCBS, Kolkata for the supercomputing facility. We thank Dr. Debashree Ghosh and Dr. Surya Kanta Ghosh for several insightful discussions and valuable suggestions on the manuscript.

## Data Availability

The data that support the findings of this study are available from the corresponding author upon reasonable request.

