## Supplementary Information for "Structural and dynamical heterogeneity of water trapped inside Na^+^-pumping KR2 rhodopsin in the dark state"

Mantu Santra[†], Aniruddha Seal[‡], Kankana Bhattacharjee[¶] and Suman Chakrabarty[*,¶]
[†] *School of Chemical and Materials Sciences, Indian Institute of Technology Goa, Ponda, Goa 403401, India*
[‡] *School of Chemical Sciences, National Institute of Science Education and Research, Bhubaneswar 752050, Odisha, India*
[¶]*Department of Chemical, Biological & Macro-Molecular Sciences, S. N. Bose National Centre for Basic Sciences, Kolkata 700106, West Bengal, India*[*]
(Dated: April 12, 2021)

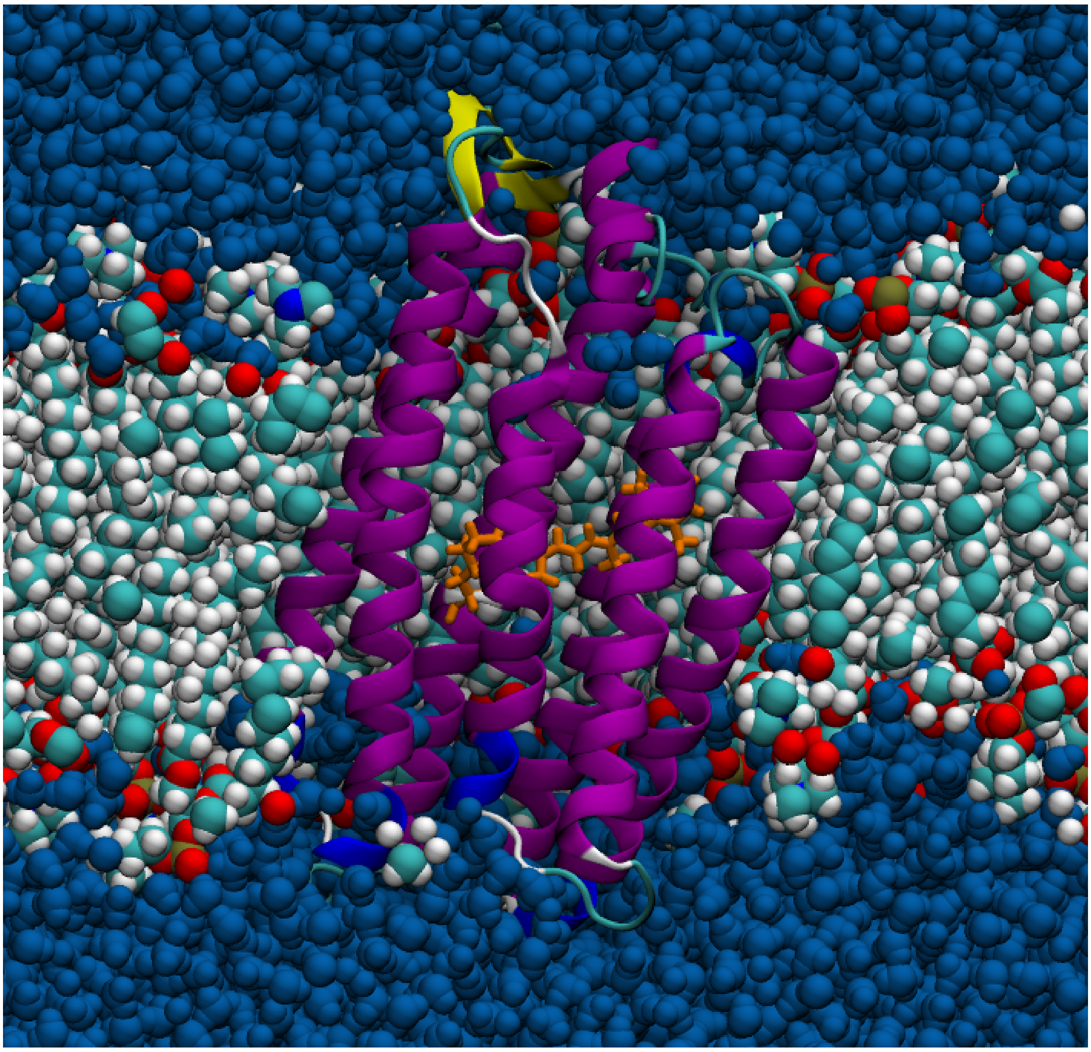

FIG. S1: **Simulated system:** The structure of KR2 Rhodopsin in the resting state embedded (purple) in a DMPC bilayer (white-cyan) with the retinal chromophore being covalently attached to the LYS255 residue (highlighted in dark yellow) at the middle of the channel . The retinal is in all-trans conformation here. The protein-membrane system is solvated with water (blue).

[*] Electronic address:

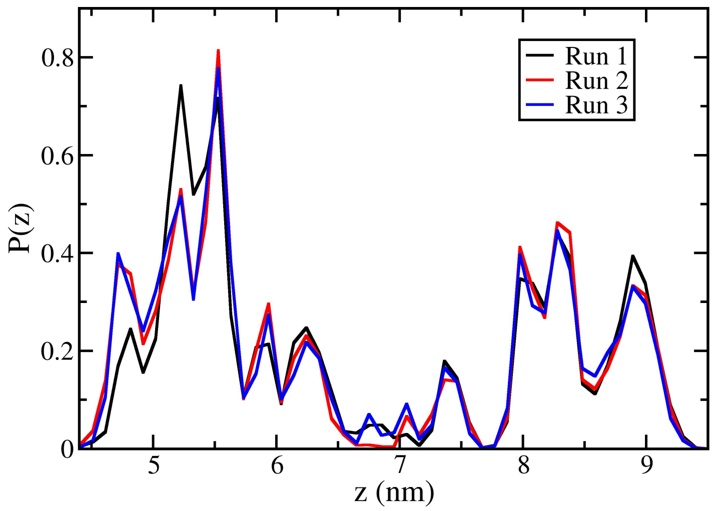


FIG. S2: The population densities of water along the $z$-direction (normal to the plane of the membrane) for three independent MD runs of length $1.5 \mu s$ each confirm reproducibility of the result in Fig.1 in the main text.

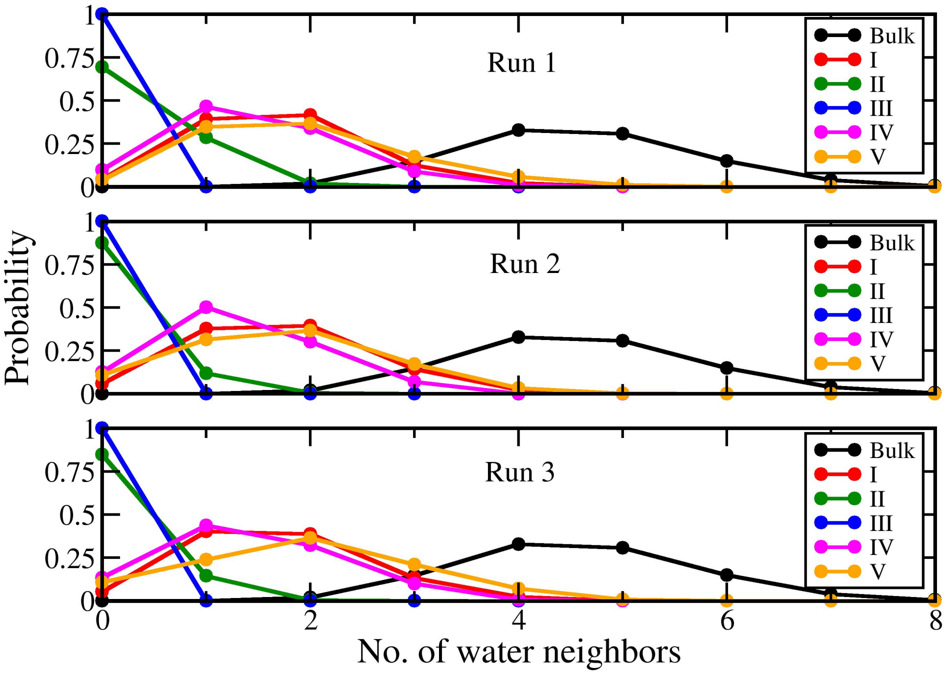


FIG. S3: The probability distributions of the number of neighboring water molecules in the first hydration shell (radius 3 Å) of the trapped water molecules for three independent MD runs of length $1.5 \mu s$ each.

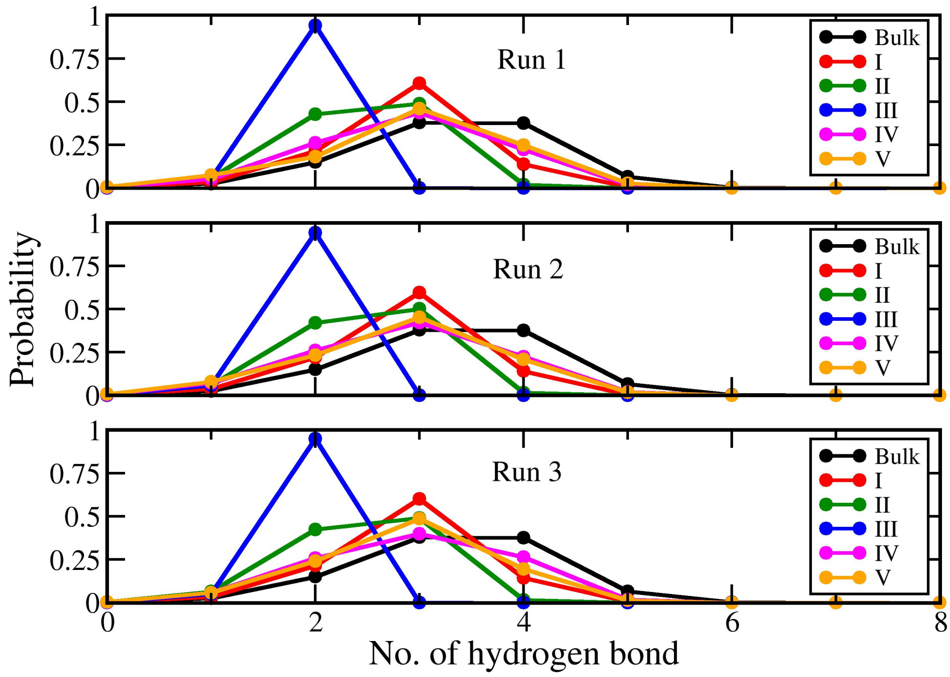


FIG. S4: The probability distributions of the number of hydrogen bond for the trapped water molecules for three independent MD runs of length 1.5$\mu s$ each.

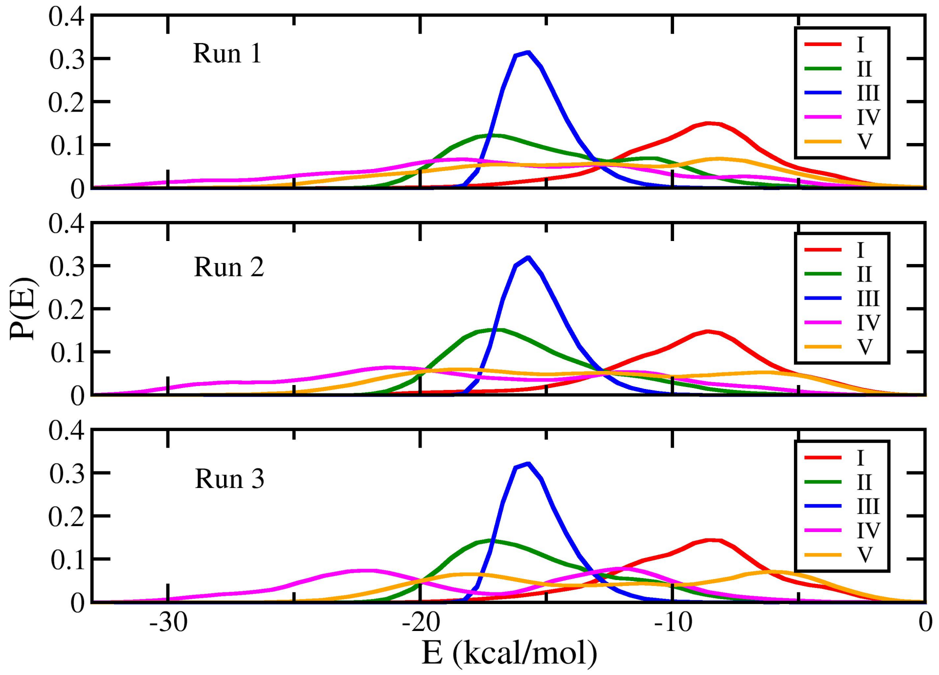


FIG. S5: The probability distributions of the water-protein interaction energy for the trapped water molecules for three independent MD runs of length 1.5$\mu s$ each.

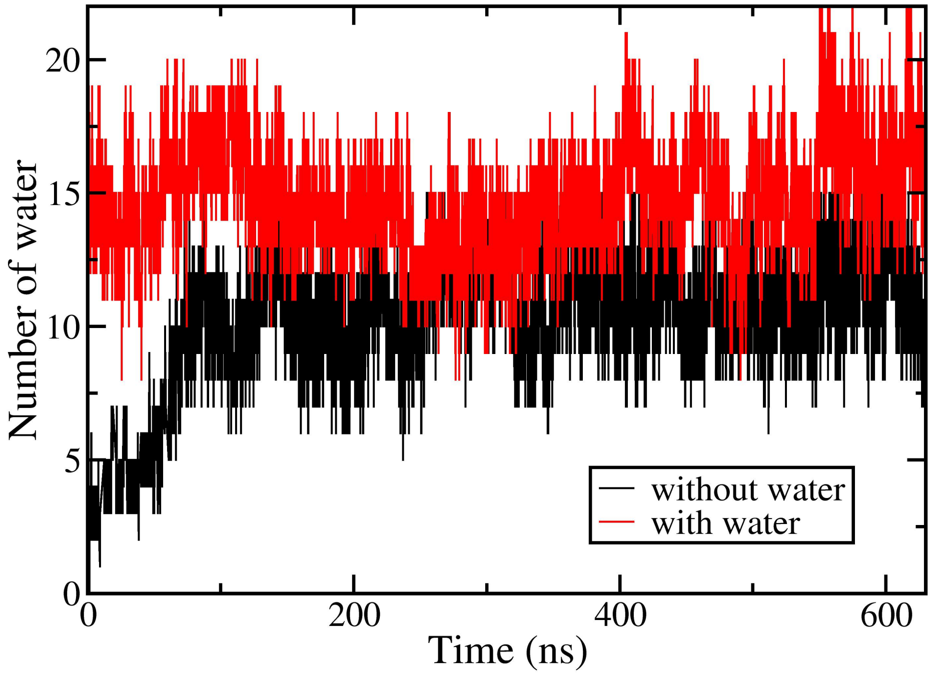


FIG. S6: The time dependence of the total number of water molecules inside KR2 channel. The black line indicates the trajectory in which all the water molecules inside the channel was removed from the initial configuration. The red line corresponds to the trajectory where the initial configuration contains water molecules inside the channel.

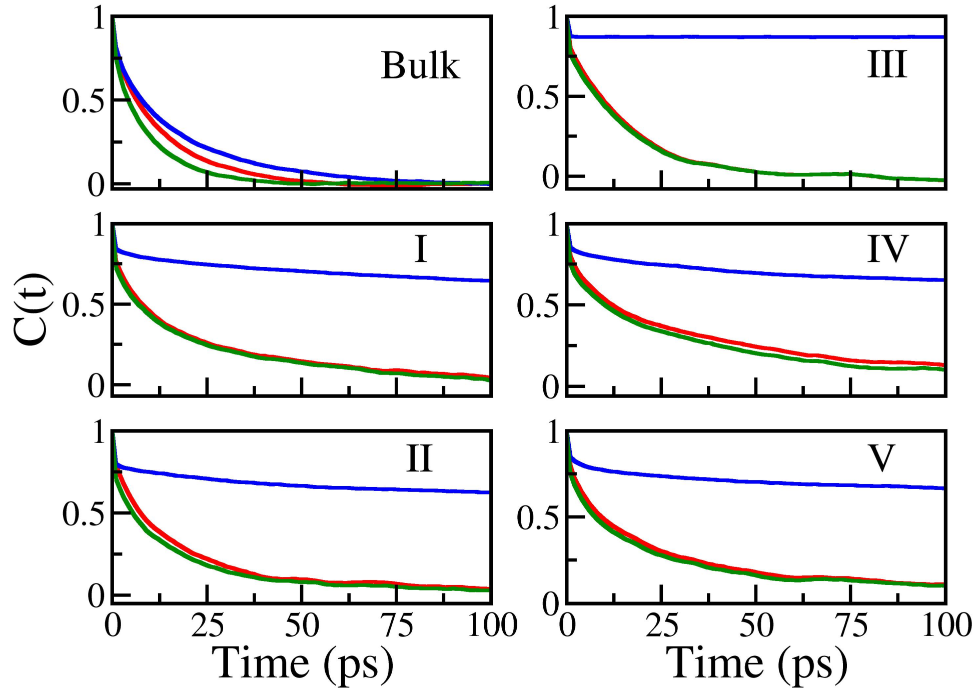


FIG. S7: Rotational time correlation functions of trapped water molecules inside the cavities including the water in the bulk for three mutually orthogonal vectors. The black, red and green lines represent rotational correlation function of the vectors corresponding to H-H bond, dipole moment of the molecule and the perpendicular direction to the plane of the molecule, respectively.
